# Correcting for physical distortions in visual stimuli improves reproducibility in zebrafish neuroscience

**DOI:** 10.1101/852962

**Authors:** Timothy W. Dunn, James E. Fitzgerald

## Abstract

Breakthrough technologies for monitoring and manipulating single-neuron activity provide unprecedented opportunities for whole-brain neuroscience in larval zebrafish^1–9^. Understanding the neural mechanisms of visually guided behavior also requires precise stimulus control, but little prior research has accounted for physical distortions that result from refraction and reflection at an air-water interface that usually separates the projected stimulus from the fish^10–12^. Here we provide a computational tool that transforms between projected and received stimuli in order to detect and control these distortions. The tool considers the most commonly encountered interface geometry, and we show that this and other common configurations produce stereotyped distortions. By correcting these distortions, we reduced discrepancies in the literature concerning stimuli that evoke escape behavior^13,14^, and we expect this tool will help reconcile other confusing aspects of the literature. This tool also aids experimental design, and we illustrate the dangers that uncorrected stimuli pose to receptive field mapping experiments.

---

In a typical zebrafish visual neuroscience experiment, an animal in water gazes at stimuli on a screen separated from the water by a small (∼500 µm) region of air (**Fig. 1a, top**). When light traveling from the screen reaches the air-water interface, it is refracted according to Snell’s law^15^ (**Fig. 1a, bottom**). At flat interfaces, a common configuration used in the literature^1,2,13^, this refraction compresses light from the screen, thereby translating and distorting the images that reach the fish (black vs. brown arrows in **Fig. 1a, bottom**). By solving Snell’s equations for this arena configuration (**Appendix 1**), we determined the apparent position of a point on the screen, *θ*, as a function of its true position, *θ*′ (**Fig. 1b**). Snell’s law implies that distant stimuli appear to the fish at the asymptotic value of *θ*(*θ*′) (∼48.6°). This implies that the entire horizon is compressed into a 97.2° “Snell window” whose size does not depend on the distances between the fish and the interface (*d*_*w*_) or the screen and the interface (*d*_*a*_), but the distance ratio *d*_*a*_/*d*_*w*_ determines the abruptness of the *θ*(*θ*′) transformation. We also calculated the total light transmittance according to the Fresnel equations (**Fig. 1b, right**). These two effects have a profound impact on visual stimuli (**Fig. 1c**). The plastic dish that contains the water has little impact on these effects (**Appendix 1**).

**Figure 1 Legend.**
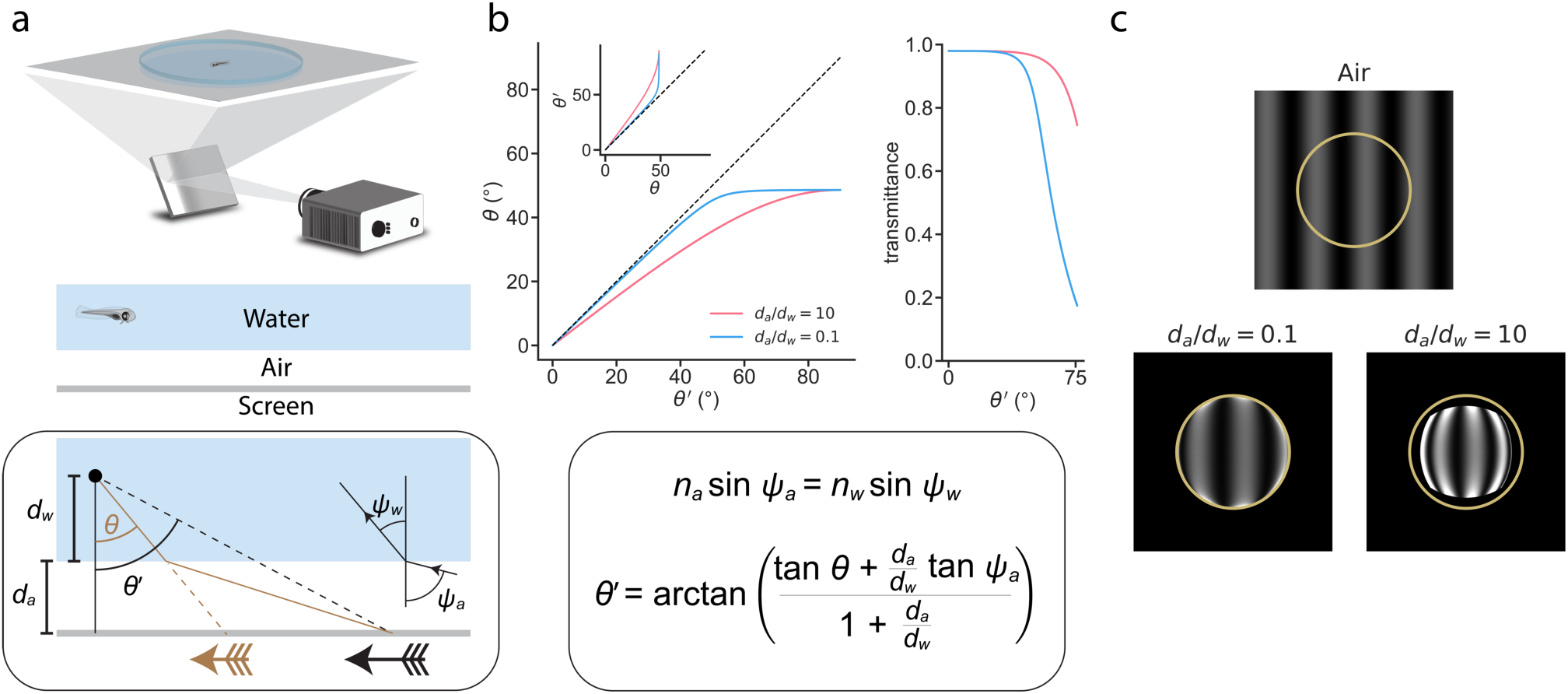
a) *Top*, In a typical zebrafish neuroscience experiment, an image is presented via projection onto a screen underneath an animal swimming in a water-filled plastic dish. *Middle*, A small layer of air separates the screen from the dish and water. Note that the effects of the plastic dish are typically minor (**Appendix 1**). *Bottom box*, This configuration causes the image received at the eye (brown arrow) to be distorted and translated relative to the projected image (black arrow). We can describe this transformation as a relationship between the true position of a projected point (*θ’*) and its apparent position (*θ*), depending on the ratio between the distance from the air-water interface to the screen (*d*_*a*_) and the distance from the eye to the air-water interface (*d*_*w*_). To solve the transformation, we use Snell’s law (illustrated in panel *b*), which relates the angle at which a light ray leaves the air-water interface (*ψ*_*w*_) to the angle at which it hits the interface (*ψ*_*a*_), depending on the refractive indices of the media (air, *n*_*a*_ = 1; water, *n*_*w*_ = 1.333). b) *Top left*, the apparent position of a point (*θ*) as a function of its true position (*θ’*), for *d*_*a*_/*d*_*w*_ = 10 (pink) and *d*_*a*_/*d*_*w*_ = 0.1 (blue), and its inverse (*inset*). *Top right*, fraction of light transmitted into the water as a function of *θ′* for the same two values of *d*_*a*_/*d*_*w*_. *Bottom box*, Using Snell’s law, we derived *θ’*(*θ*) (*top left inset*), whose inverse we take numerically to arrive at *θ*(*θ’*) (*top left*). c) Simulated distortion of a standard sinusoidal grating. Yellow circle denotes the extent of the Snell window (∼97.2° visual angle). Contrast axes are matched across panels and saturate to de-emphasize the ring of light at the Snell window, whose magnitude would be attenuated by unmodeled optics in the fish eye (**Materials and methods**).

These distortions have the potential to affect quantitative conclusions drawn from studies of sensorimotor transformations. For example, Temizer et al.^14^ and Dunn et al.^13^ both found that a critical image size triggered looming-evoked escape behavior, but they reported surprisingly different values for the critical angle (21.7° ± 4.9° and 72.0° ± 2.5°, respectively, mean ± 95% CI). This angular discrepancy of 50.3° is about 14% of the maximal possible angle (360°). A major difference in experiment design is that Temizer et al. showed stimuli from the front through a curved air-water interface, and Dunn et al. showed stimuli from below through a flat air-water interface (**Fig. 2a**). Correcting with Snell’s law, and quantifying the size of irregularly shaped stimuli with their solid angle, suggests that the Dunn et al. stimulus spanned just 0.24 steradians, or 1.9% of the maximal angular size (4π steradians), when the fish exhibited its escape response, rather than the uncorrected 20% (**Fig. 2b, Appendix 1, Materials and methods, Supplementary Movie**). The same correction applied to Temizer et al. reduces the critical size to 0.6% from 6% of the maximal angular size (**Fig. 2b, Appendix 2, Materials and methods**). Correcting with Snell’s law thus markedly reduced this discrepancy in the literature, shrinking an original discrepancy of 14.0% ± 1.6% to 1.3% [+0.6, -0.5]% (**Fig. 2c**), a result not explained by the solid angle conversion alone (7.2% [+1.2, -1.0]% discrepancy). The small remaining difference could indicate an ethologically interesting dependence of behavior on the spatial location of the looming stimulus^13,14^.

**Figure 2 Legend.**
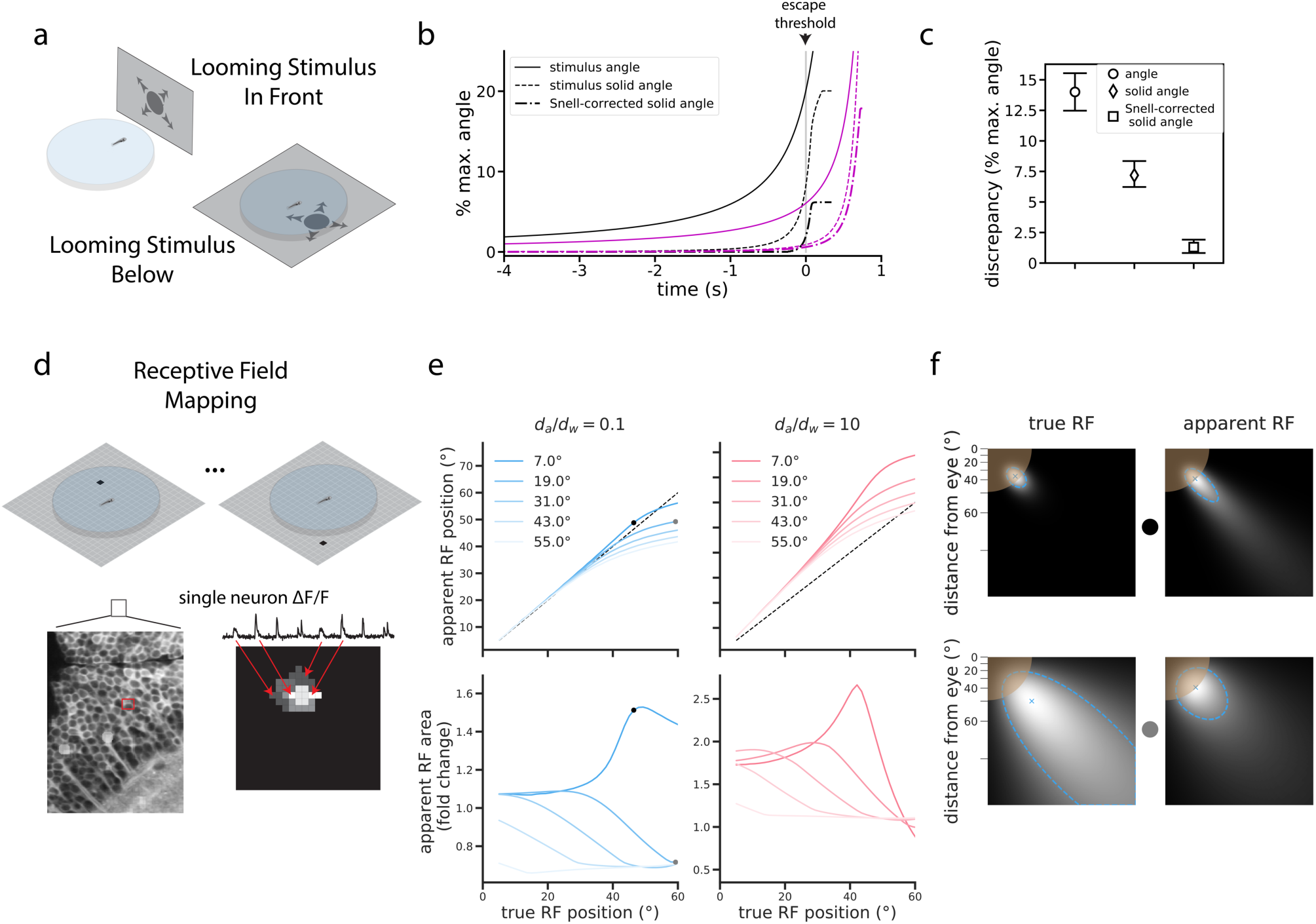
a) In the zebrafish literature, two configurations were used to probe the neural circuitry processing looming stimuli that expand over time. In one, fish were embedded off-center in a curved plastic dish and a screen presented stimuli in front of the animal through the curved interface of the dish (Temizer et al.). In the other, fish were embedded (or swam freely) in a similar dish, but stimuli were presented on a screen below the dish (as in panel *a*) (Dunn et al.). b) Plot detailing the changes to the looming expansion time courses after correcting for Snell’s law and converting to solid angle, which more accurately describes the irregular stimulus shapes produced by the optical distortion (**Materials and methods**). Percentages indicate fractional values, relative to the maximal stimulus angle (2π radians) and maximal stimulus solid angle (4π steradians). Curves corresponding to Dunn et al. and Temizer et al. are plotted in black and magenta, respectively. c) Snell’s law corrections reduced the discrepancy between Dunn et al. and Temizer et al. d) In a simple receptive field (RF) mapping experiment, dots appear at different positions on a screen (*Top*), and behavioral or neural responses (*Bottom*) are measured. In the latter case, a map of a single neuron’s RF is constructed by assigning the measured *ΔF*/*F* signal to the point on the screen that evoked the *ΔF*/*F* response. e) Snell’s law predicts changes in RF peak positions (*Top*) and RF sizes (*Bottom*). The magnitude of these changes depends on the true RF position, size, and *d*_*a*_/*d*_*w*_. The black dots indicate the RFs in *panel f, top*, and the gray dots show the RFs in *panel f, bottom*. f) Illustrations of two simulated “true” RFs and their corresponding measurement distortions predicted using Snell’s law. For simplicity, we show only one quadrant of the screen space, with the fish at the top left corner. The brown circle denotes the extent of the Snell window. As RFs are mapped directly to screen pixels, the axes are non-linear in terms of angle relative to the fish (top left corner). Each blue “x” denotes the peak position of the RF displayed in each plot. The dashed blue border denotes the half-maximum value of each RF, and the size of the RF is the area within one of these borders.

Accounting for optical distortions will be critical for understanding other fundamental properties of the zebrafish visual system. For example, a basic property of many visual neurons is that they respond strongest to stimuli presented in one specific region of the visual field, termed their receptive field (RF)^11^. When we simulated the effect of Snell’s law on RF mapping under typical experimental conditions (**Fig. 2d**), we predicted substantial errors in both the position and size of naively measured receptive fields (**Fig. 2e, Methods**). Depending on the properties of the true RF, its position and size could be either over- or under-estimated (**Figs. 2e-f**), with the most drastic errors occurring for small RFs appearing near the edge of the Snell window.

Future experiments could avoid distortions altogether by adjusting experimental hardware. For instance, fish could be immobilized in the center of water-filled spheres^11,16^, or air interfaces could be removed altogether, such as by placing a projection screen inside the water-filled arena. But in practice the former would restrict naturalistic behavior, and the latter would reduce light diffusion by shrinking the refractive index mismatch necessary to transmit stimuli over a large range of angles. An engineering solution might build diffusive elements into the body of the fish tank^10^. Alternatively, we propose a simple computational solution to account for expected distortions when designing stimuli or analyzing data. Our tool (https://www.github.com/spoonsso/snell_tool/) converts between normal and distorted image representations for the most common zebrafish experiment configuration (**Fig. 1a**), and other geometries could be analyzed similarly. This tool will therefore improve the interpretability and reproducibility of innovative experiments that capitalize on the unique experimental capabilities available in zebrafish neuroscience.

## Acknowledgments

We thank Damon Clark, Ruben Portugues, and Kristen Severi for helpful comments on the manuscript. We also thank Florian Engert and Haim Sompolinsky for early support and partial funding of the project (NIH grant U01 NS090449). TWD was supported by Duke Forge and Duke AI Health. JEF was supported by the Howard Hughes Medical Institute.

## Materials and methods

See **Appendix 1** and **Appendix 2** for the geometric consequences of Snell’s Law at flat and curved interfaces, respectively.

### Implications of the Fresnel equations

Only a portion of the incident light is transmitted into the water to reach the eye. We calculated the fraction of transmitted light according to the Fresnel equations. Assuming the light is unpolarized,

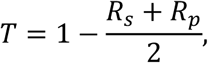

where *T* is the fraction of light transmitted across an air-water interface at incident angle ψ_*a*_ = ψ_*a*_(*θ*) (See **Appendices 1, 2**), and

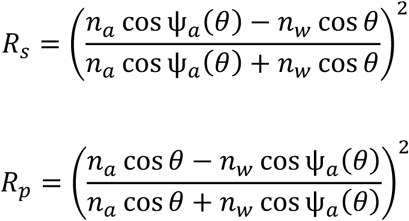

are the reflectances for s-polarized (i.e. perpendicular) and p-polarized (i.e. parallel) light, respectively. When including the plastic dish in our simulations, we modified these equations to separately calculate the transmission fractions across the air-plastic and the plastic-water interfaces. We assumed that the full transmission fraction is the product of these two factors, thereby ignoring the possibility of multiple reflections within the plastic.

### Illustrating distorted sinusoidal gratings

For all image simulations in **Fig. 1c**, we neglected the plastic and fixed the total distance between the fish and the virtual screen, *d*_*a*_ + *d*_*w*_, to be 1 cm, a typical distance in real-world experiments. The virtual screen was considered to be a 4 × 4 cm square with 250 pixels / cm resolution. To transform images on the virtual screen, we shifted each light ray (i.e. image pixel) according to Snell’s Law, scaled its intensity according to the Fresnel equations, and added the intensity value to a bin at the resulting apparent position. This simple model treats the fish eye as a pinhole detector, whereas real photoreceptors blur visual signals on a spatial scale determined by their receptive field. Consequently, our simulation compresses a large amount of light onto the overly thin border of the Snell window, and we saturated the grayscale color axes in **Fig. 1c** to avoid this visually distracting artifact.

To make the image as realistic as possible, we mimicked real projector conditions using gamma-encoded gratings with spatial frequency 1 cycle / cm, such that

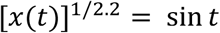

with *x*(*t*) ranging from 1.0 to 500.0 lux, a standard range of physical illuminance for a lab projector. The exponent on the left represents a typical display gamma encoding with gamma = 2.2. To reduce moiré artifacts arising from ray tracing, we used a combination of ray supersampling (averaging the rays emanating from 16 sub-pixels for each virtual screen pixel) and stochastic sampling (the position of each ray was randomly jittered between [-1 1] from its native position) ^17^. In **Fig. 1c**, we display the result of these operations followed by a gamma compression to mimic the perceptual encoding of the presented stimulus.

### Corrections to looming visual stimuli

We approximated the geometric parameters from Dunn et al. (flat air-water interface, *d*_*a*_ = 0.5 mm, *d*_*w*_ = 3 mm, *d*_*p*_ = 1 mm, stimulus offset from the fish by 10 mm along the screen) and Temizer et al. (curved air-water interface, *d*_*a*_ = 8 mm, *d*_*w*_ = 2 mm, *d*_*p*_ = 1 mm, r = 17.5 mm, stimulus centered) to create Snell-transformed images of circular stimuli with sizes growing over time (**Figs. 2a-c**). We used a refractive index of *n*_*p*_ = 1.55 for the polystyrene plastic. While Dunn et al. collected data from freely swimming fish, the height of the water was kept at 5 mm, and 3 mm reflects a typical swim depth. Since freely swimming zebrafish can adjust their depth in water, it’s an approximation to treat *d*_*w*_ as constant.

We quantified the size of each transformed stimulus with its solid angle, the surface area of the stimulus shape projected onto the unit sphere. To calculate the solid angle, we first represented stimulus border pixels in a spherical coordinate system locating the fish at the origin. The radial coordinate does not affect the solid angle, so we described each border pixel by two angles: the latitude, *α*, and longitude, *β*. To calculate the area, we used an equal-area sinusoidal (Mercator) projection given by

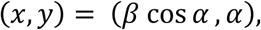

which projects an arbitrary shape on the surface of a sphere onto the Cartesian plane. While distances and shapes are not conserved in this projection, area as a fraction of the sphere’s surface area is maintained. Thus, we could calculate the solid area of the stimulus in this projection by finding the area of the projected 2D polygon.

### Receptive field mapping

We simulated receptive field (RF) mapping experiments by tracing light paths from single pixels on a virtual screen to the fish (**Figs. 2d-f**). We modeled a neuron’s RF as a Gaussian function on the sphere, defined the “true RF” to be the pixel-wise response pattern that would occur in the absence of the air-water interface, and defined the “apparent RF” as the pixel-wise response pattern that would be induced with light that bends according to Snell’s law at an air-water interface. More precisely, we modeled the neural response to pixel activation at position *x* as

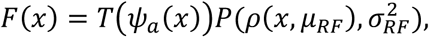

where *T*(*ψ*_*a*_(*x*) is the fraction of light transmitted (Fresnel equations), *μ*_*RF*_ and *σ*_*RF*_ are the mean and standard deviation of the Gaussian RF, *ρ*(*x, μ*_*RF*_) is the distance along a great circle from the center of the RF to the pixel’s projected retinal location, and 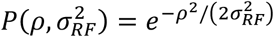 is the Gaussian RF shape. We calculated the great circle distance between points on the sphere as

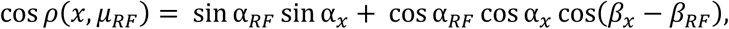

where (*α*_*RF*_, *β*_*RF*_) are the latitude and longitude coordinate of the RF center, and (*α*_*x*_, *β*_*x*_) are the latitude and longitude coordinates of the projected pixel location. We quantified the position of the RF as the maximum of *F*(*x*), converted to an angular coordinate along the screen. We quantified RF area as the solid angle of the shape formed by thresholding *F*(*x*) at half its maximal value.

## Appendix 1

### Implications of Snell’s Law at a flat interface

For this and all subsequent analyses, we treat the fish as a pinhole detector. Here we derive *θ*′(*θ*) with the aid of **Appendix Fig. 1**. Note that this derivation includes optical effects from the plastic dish, but these effects will be relatively minor. To begin, we summarize the basic trigonometry of the problem. The true angular position of the stimulus is given by

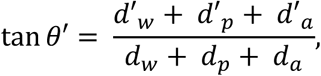

where *d*_*w*_ is the normal distance between the fish and the water-plastic interface, *d*_*p*_ is the normal distance between the water-plastic and plastic-air interfaces, *d*_*a*_ is the normal distance between the air interface and the screen (interface and screen assumed to be parallel), *d*′_*w*_ is the parallel distance traveled by the light ray in the water, *d*′_*p*_ is the parallel distance traveled by the light ray in the plastic, and *d*′_*a*_ is the parallel distance traveled by the light ray in air. Each parallel distance is related to the corresponding normal distance by simple trigonometry. The apparent angular location of the stimulus satisfies

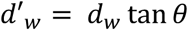

and the incident light angle satisfies

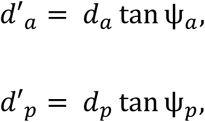

thereby leading to

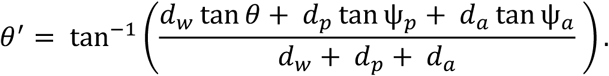

We can next use Snell’s Law to reduce the number of angular variables. In particular,

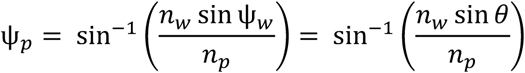

and

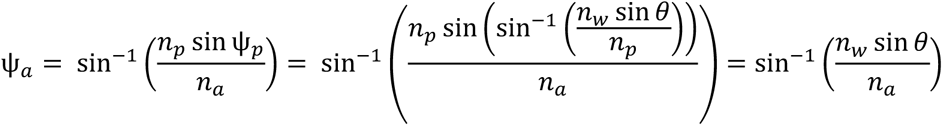

together imply that

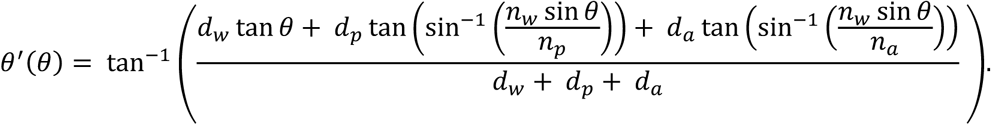

The role of plastic in this equation is typically minimal. To see this, first note that *n*_*w*_ ≈ 1.333 < *n*_*p*_ ≈ 1.55, which implies that 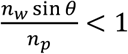. This implies that the Snell window is determined by 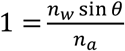, and the properties of the high-index plastic dishes have no effect on the size of the Snell window. The plastic can cause distortions within the Snell window, but these effects were small for all experimental arenas analyzed in this paper, as we empirically found that none of our results qualitatively depended upon the plastic. We therefore chose to highlight the critical impact of the air-water interface by assuming that *d*_*p*_ = 0 in the main text’s conceptual discussion. We nevertheless included nonzero values of *d*_*p*_ in our computational tool so that users can account for the quantitative effects of the plastic dish. We also included the effects of plastic when quantitatively correcting previously published results. Because analytically inverting *θ*′(*θ*) is non-trivial, we noted from the graph of *θ*′(*θ*) that the inverse function exists and calculated *θ*(*θ*′)with a numerical look-up table (*e.g.* **Fig. 1b**).

**Appendix Figure 1 Legend.**
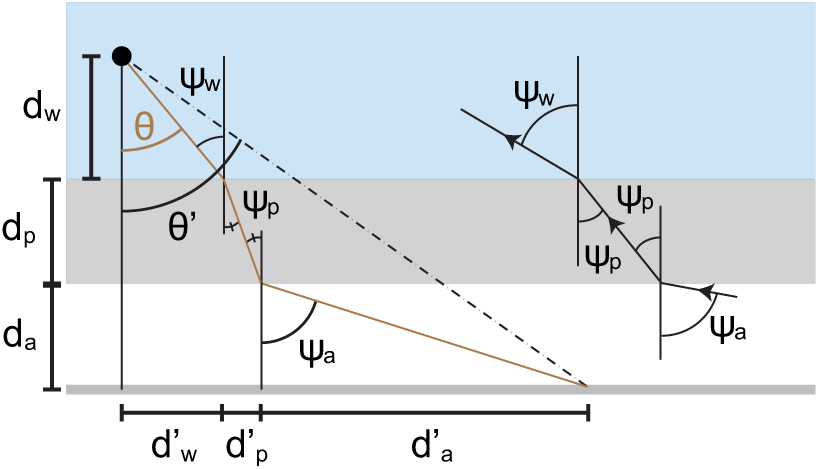
Illustration of mathematical variables used to analyze optical distortions in arena geometries where flat air-plastic and plastic-water interfaces separate the fish from the projection screen. The brown line denotes the trajectory of a light ray traveling from the screen to the fish. We quantify the image transformation by relating the true angular position of each projected point (*θ’*) to its apparent position (*θ*). The derivation involves several distances (e.g. *d*_*w*_) that summarize the ray’s trajectory through air (white region), plastic (gray region), and water (blue region). Refraction angles (*ψ*_*a*_, *ψ*_*p*_, *ψ*_*w*_) describe the bending of light at each interface.

## Appendix 2

### Implications of Snell’s Law at a curved interface

When the fish is mounted off-center (**Appendix Fig. 2a)** in a circular dish (brown dot), rays pass through a curved interface and are refracted at tangent lines. We begin by using Snell’s law and basic trigonometry to relate each refraction angle to *θ*. Let *d*_*a*_ denote the distance in air between the edge of the plastic dish and the screen, *d*_*p*_ denote the thickness of the plastic dish, *d*_*w*_ denote the distance in water between the fish and the edge of the tank nearest the screen, and *r* denote the radius of the dish (excluding the plastic). We assume that *d*_*w*_ ≤ *r* and the screen is perpendicular to the line between the fish and the center of the dish. Cases where the fish is behind the dish’s center or the screen is angled can be analyzed similarly. Starting at the fish and moving outwards, we first apply the Law of Sines to the gray triangle to find

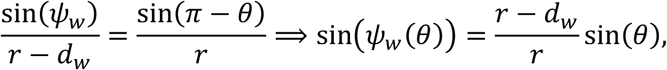

where we’ve used the identity sin(*π* − *x*) = sin(*x*). It will be useful for later to note that this triangle also implies that *γ* = *π* – (*ψ*_*w*_ + *π* − *θ*) = *θ* − *ψ*_*w*_. Snell’s Law at the plastic-water interface implies,

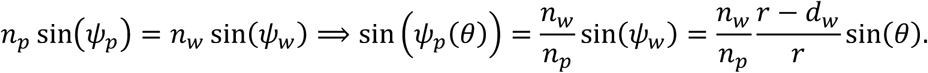

We next relate the two plastic refraction angles to each other by applying the Law of Sines to the orange triangle and find

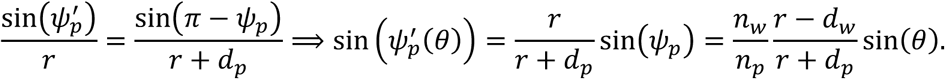

Finally, we determine the dependence of ψ_*a*_ on *θ* from Snell’s Law applied to the air-plastic interface,

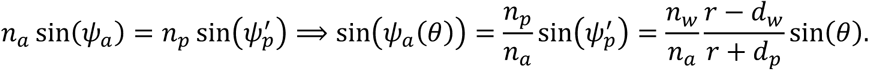

With these formulae in hand, we now proceed to the main goal of deriving an expression for *θ*′(*θ*). Since we’ve already extracted everything from Snell’s Law, all that remains is basic trigonometry, which we illustrate in **Appendix Fig. 2b**. First note that applying the definition of the tangent function to the blue triangle implies that

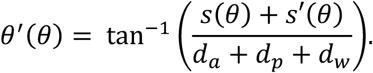

It thus suffices to determine expressions for *s*(*θ*) and *s*′(*θ*). Consider first *s*′(*θ*). The large red triangle implies

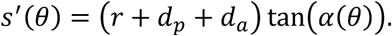

Rewriting *α* in terms of the other two angles in the 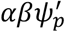 triangle gives 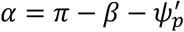. Rewriting *β* in terms of the other two angles in the *βγψ*_*p*_ triangle gives *β* = *π* – (*γ* + *ψ*_*p*_) = *π* – *θ* + *ψ*_*w*_ – *ψ*_*p*_. Putting these pieces together, we thus find

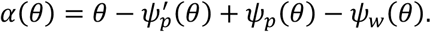

Next consider *s*(*θ*). Applying the Law of Sines to the green triangle, we find

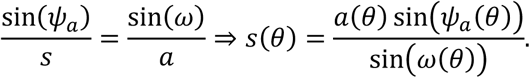

Rewriting *ω* in terms of the other two angles in the green triangle, *ω* = *π* – (*ψ*_*a*_ + *π* − *φ*) = *φ* − *ψ*_*a*_, and rewriting *φ* in terms of the other angles in the red triangle, 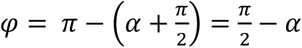, we find

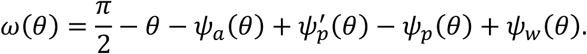

Finally, we find the dependence of *a* on *θ* from the red triangle using the definition of the cosine function

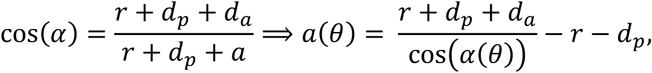

Since we’ve written *α, α, ω*, and the refraction angles as functions of *θ*, we’ve fully specified *s*(*θ*), *s*′(*θ*), and *θ*′(*θ*). As with the flat interface, we calculated *θ*(*θ*′) using a numerical look-up table.

**Appendix Figure 2 Legend.**
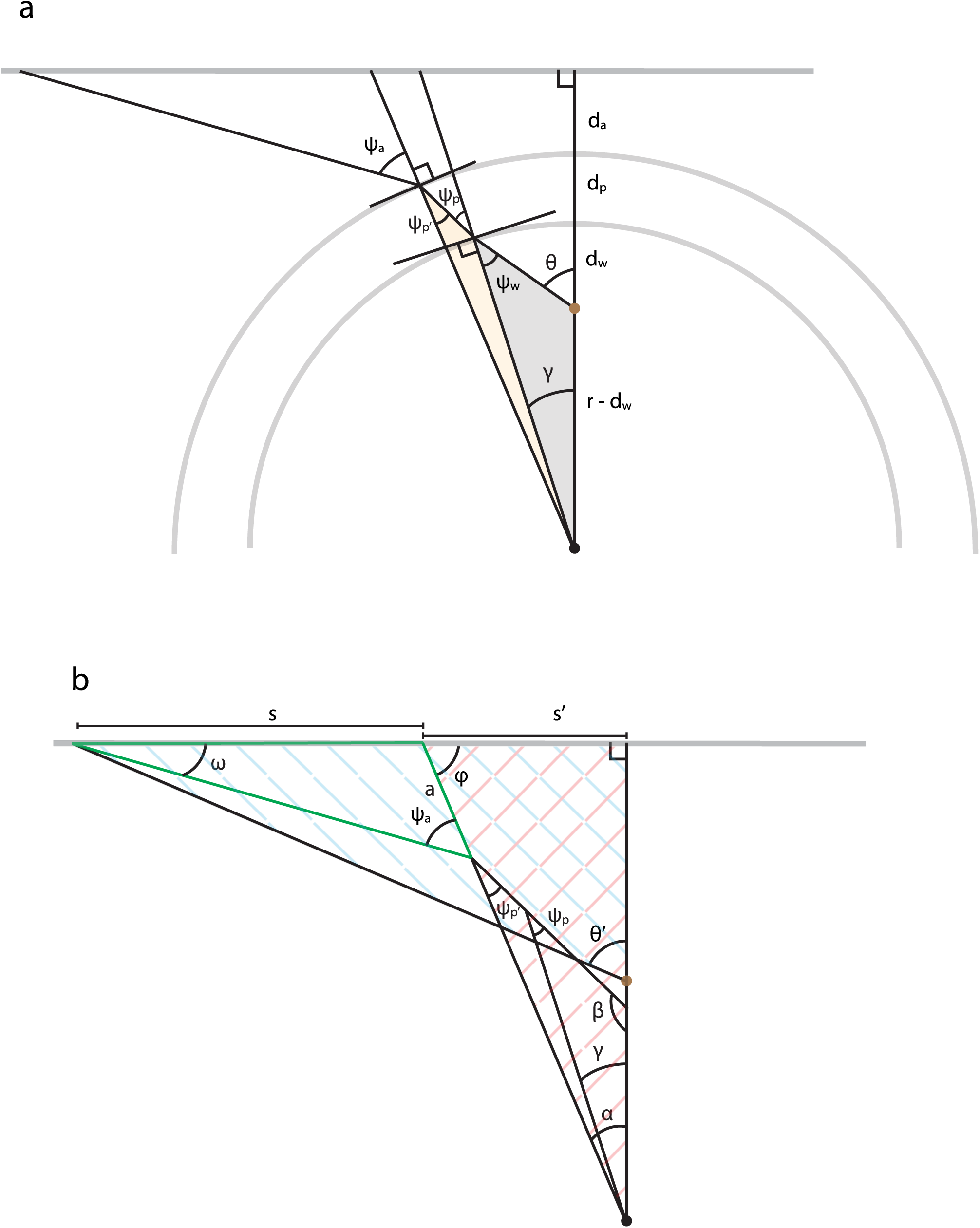
a) Illustration of mathematical variables used to analyze optical refraction in arena geometries where curved air-plastic and plastic-water interfaces separate the fish (brown dot) from the projection screen. We assume that the interfaces are circular, that the fish is mounted off-center, and that the screen and fish are at the same elevation. The last assumption avoids distortions that could result from the flat vertical interface running parallel to the longitudinal axis of the cylindrical dish. We denote the radius of the arena’s water-filled compartment as *r*. The derivation additionally involves several distances that summarize the placement of the fish in the dish (*d*_*w*_), the thickness of the plastic (*d*_*p*_), and the distance separating the dish from the screen (*d*_*a*_). Refraction angles 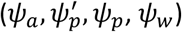 are relative to each interface’s normal vector and describe the bending of light. Each shaded region highlights a triangle whose trigonometry is helpful for relating the refraction angles to the apparent angular position of a light source (*θ*). b) Illustration of mathematical variables used to trigonometrically relate the true angular position of each projected point (*θ’*) to its apparent position (*θ*), assuming the same arena geometry as panel *b*. The derivation utilizes most triangles shown, several of which are cross-hatched or outlined to direct the reader’s eye.

## References

1. Ahrens, M. B. et al. Brain-wide neuronal dynamics during motor adaptation in zebrafish. Nature (2012). doi:10.1038/nature11057

2. Vladimirov, N. et al. Light-sheet functional imaging in fictively behaving zebrafish. Nat. Methods 11, 1–2 (2014).

3. Prevedel, R. et al. Simultaneous whole-animal 3D imaging of neuronal activity using light-field microscopy. Nat. Methods 11, 727–730 (2014).

4. Kim, D. H. et al. Pan-neuronal calcium imaging with cellular resolution in freely swimming zebrafish. Nat. Methods 14, 1107–1114 (2017).

5. Naumann, E. A. et al. From Whole-Brain Data to Functional Circuit Models: The Zebrafish Optomotor Response. Cell 167, 947-960.e20 (2016).

6. Ahrens, M. B., Orger, M. B., Robson, D. N., Li, J. M. & Keller, P. J. Whole-brain functional imaging at cellular resolution using light-sheet microscopy. Nat. Methods 10, (2013).

7. Portugues, R., Feierstein, C. E., Engert, F. & Orger, M. B. Whole-brain activity maps reveal stereotyped, distributed networks for visuomotor behavior. Neuron 81, 1328–43 (2014).

8. Dunn, T. W. et al. Brain-wide mapping of neural activity controlling zebrafish exploratory locomotion. Elife (2016). doi:10.7554/eLife.12741

9. Vladimirov, N. et al. Brain-wide circuit interrogation at the cellular level guided by online analysis of neuronal function. Nat. Methods 15, 1117–1125 (2018).

10. Stowers, J. R. et al. Virtual reality for freely moving animals. Nat. Methods 14, 995–1002 (2017).

11. Zhang, Y. & Arrenberg, A. B. High throughput, rapid receptive field estimation for global motion sensitive neurons using a contiguous motion noise stimulus. J. Neurosci. Methods 326, 108366 (2019).

12. Sajovic, P. & Levinthal, C. Inhibitory mechanism in zebrafish optic tectum: visual response properties of tectal cells altered by picrotoxin and bicuculline. Brain Res. 271, 227–40 (1983).

13. Dunn, T. W. et al. Neural Circuits Underlying Visually Evoked Escapes in Larval Zebrafish. Neuron 89, 1–16 (2016).

14. Temizer, I., Donovan, J. C., Baier, H. & Semmelhack, J. L. A Visual Pathway for Looming-Evoked Escape in Larval Zebrafish. Curr. Biol. 25, 1823–1834 (2015).

15. Hecht, E. Optics. (Pearson Education, 2016).

16. Dehmelt, F. A. et al. Spherical arena reveals optokinetic response tuning to stimulus location, size and frequency across entire visual field of larval zebrafish. bioRxiv 754408 (2019). doi:10.1101/754408

17. Dippé, M. A. Z. & Wold, E. H. Antialiasing Through Stochastic Sampling. SIGGRAPH Comput. Graph. 19, 69–78 (1985).

